# Troponin-I localizes selected apico-basal cell polarity signals

**DOI:** 10.1101/404863

**Authors:** Sergio Casas-Tintó, Alberto Ferrús

## Abstract

Beyond its well characterized role in muscle contraction, *Drosophila* Troponin I (TnI) is expressed in other cell types where it plays a role in proliferation control. TnI traffics between the nucleus and the cytoplasm through a sumoylation-dependent mechanism. We address here the role of TnI in the cytoplasm. TnI accumulates in the apical region of epidermal cells and neuroblasts. TnI helps to localize and co-immunoprecipitates with Par-3/Bazooka and with disc large (Dlg), two components of the apico-basal polarity system. By contrast, Scribbled is not affected by TnI depletion. In neuroblasts, TnI is required for the polar localization of Miranda while non-polar Dlg is not affected. TnI loss-of-function triggers genome instability, cell apoptosis and extrusion from wing disc epithelia. However, rescue from apoptosis by p35 does not prevent genome instability demonstrating that both features, apoptosis and genome instability, are mechanistically independent. While PI3K is known to contribute to apico-basal polarity of epithelia in vertebrates, *Drosophila* PI3K depletion alters neither the apical localization of TnI or Par3/Bazooka, nor the basal localization of Dlg. However, the overexpression of PI3K prevents the polarity defects caused by TnI depletion. Thus, TnI binds certain apico-basal polarity signals in a cell type dependent context, and it unveils a hitherto unsuspected diversity of mechanisms to allocate cell polarity factors.

## INTRODUCTION

Cell polarity results from the focal accumulation of proteins and organelles and is a key determinant of multiple features of cell biology from cell proliferation to organ morphogenesis (Harris and Peifer, 2005; Luxton and Gundersen, 2011; St Johnston and Ahringer, 2010). Out of the several cell polarity systems known, the apico-basal one (AB) is characteristic of epithelia. AB polarity is internal to the cell and some of its determinant protein complexes have been identified (Humbert et al., 2003; Rolls et al., 2003; Wodarz et al., 2000). AB polarity is unlikely to result from a single universal mechanism, even among epithelial tissues. For example, Bazooka/Par3 is a main requirement for AB polarity in many epithelial cells while it plays an auxiliary role, if at all, in the follicular type in the *Drosophila* ovaries (Shahab et al., 2015). In addition to cell internal factors, mechanical interactions with neighboring cells influence the establishment of membrane domains in a polarized manner (Asnacios and Hamant, 2012; Goehring and Grill, 2013). As all cellular features, cell polarity is dynamic and the corresponding factors move along mostly conserved actin, myosin II, keratin and tubulin cytoskeletal scaffolds (Flores-Benitez and Knust, 2016; Monier et al., 2015; Noordstra et al., 2016; Salas et al., 2016). Presumably, motor systems would provide the required force to execute these movements. However, how these cytoskeletal apparatuses regulate their activity and their directional traffic remains largely unknown.

The phosphoinositide-3-kinase (PI3K) is a family of lipid kinases of which class I members are most thoroughly characterized. While vertebrates have four isoforms of the class I PI3K, *Drosophila* has only one. All forms, however, signal through the generation of the phosphatidylinositol (3,4,5)-triphosphate [PtdIns(3,4,5)P3] lipid that regulates AB polarity and epithelial cell morphogenesis, among other functions (Comer and Parent, 2007; Gibson and Perrimon, 2003; Krahn and Wodarz, 2012; Shewan et al., 2011). Also, PI3K is often upregulated in several forms of cancer (Juvin et al., 2013; Tzenaki and Papakonstanti, 2013) suggesting a chain of causal links between excess of PI3K activity, loss of AB cell polarity, aberrant mitosis, genome instability and, finally, tumorous growth (McCaffrey et al., 2012). Whether these events are indeed causally linked or if, by contrast, they are mechanistically independent is still open.

We have shown previously that *Drosophila* TnI is expressed early in the syncytial embryo before muscles or any other cell types are specified. The TnI protein traffics between the nucleus and the cytoplasm as a function of cell cycle status using a sumoylation-dependent mechanism (Sahota et al., 2009). We have also shown that TnI is required for cell proliferation and synergizes with classical oncogenes *Ras, Lgl* and *N* for tumor outgrowth (Casas-Tinto et al., 2016). Here, we set out to characterize the mechanisms by which TnI depletion affects apico-basal polarity of epithelial cells in the wing imaginal discs. In addition, we address the context dependence of AB polarity mechanisms by comparing TnI depleted epithelial cells and neuroblasts, a cell type that divide by asymmetric divisions and their arrangement is not epithelial.

## Results

### TnI can show apical localization

To visualize TnI we use the validated monoclonal antibody SC1 (Casas-Tinto et al., 2016). Wing disc cells show a strong TnI signal, which is localized apically with respect to baso-lateral Scribbled (**Fig. 1A,B**). In its apical position, TnI co-localizes with Actin (**Fig. 1C-E**). This is consistent with the canonical actin-binding function of TnI. In neuroblasts, TnI is also accumulated in apical position with respect to Prospero (**Fig. 1F**). Neuroblasts divide asymmetrically to yield another neuroblast and a ganglion mother cell (GMC). This later cell type undergoes symmetric divisions to yield neuron or glial cells. TnI is transiently found in the nucleus of GMC and their daughter cells (**Fig. 1G**). In salivary gland cells, whose large size facilitates visualization of protein localization, TnI is also accumulated in the apical domain with respect to the basolateral Dlg and the dorsal Crumbs (**Fig 1H**). These large cells allow revealing that TnI accumulates in the apical domain of the cell nucleus (**Fig. 1I**). Thus, in all cell types analyzed TnI shows apical localization in the cytoplasm and, possibly, in the nucleus also.

**Figure 1.**
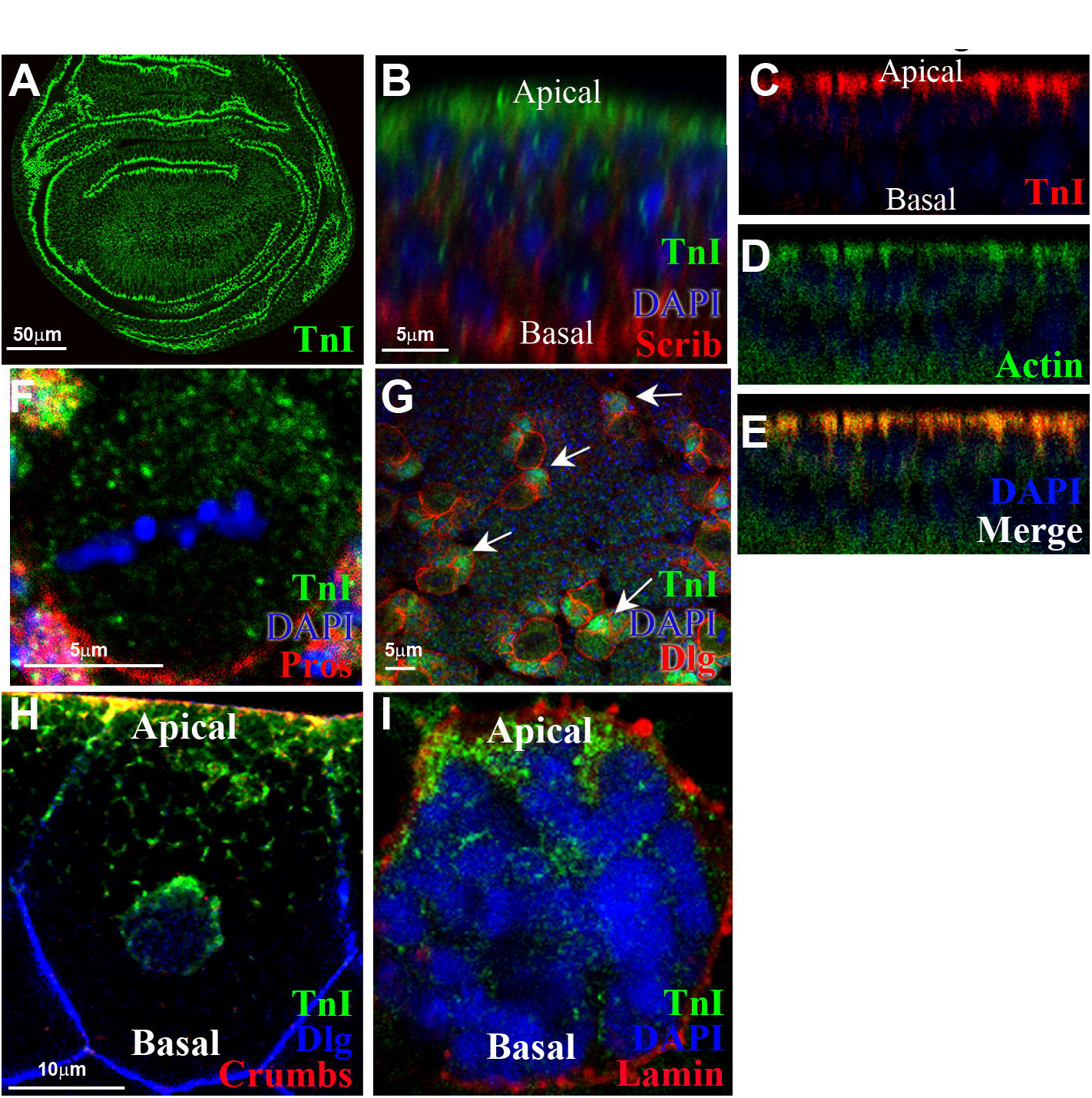
TnI can show polar localization. A) Imaginal wing disc immunostained against TnI with SC1 antibody. **B)** The Z-axis view of a wing disc shows apical TnI (green) in relation to basal Scrib (red). Nuclei are stained by DAPI (blue). **C-E)** Z-axis view of a wing disc immunostained against TnI and Actin. Note the co-localization of both proteins in the apical side of cells. **F)** Larval neuroblast showing apical TnI (green) with respect to Prospero (red). **G)** Larval brain stained for Dlg (red) and TnI (green). Note that TnI accumulates in the nucleus of ganglion mother cells (arrow) but not in differentiating brain cells where it is mostly cytoplasmic. **H)** Salivary gland cell immunostained against TnI (green), Crumbs (red) and Dlg (blue). Note the apical localization of TnI. **I)** Nucleus of a salivary gland cell stained for Lamin (red) to mark the nuclear membrane, DAPI (blue) to mark the chromatin and TnI (green).

### TnI helps to localize other apico-basal polarity proteins

The polar localization of TnI prompted an analysis of well characterized polarity proteins under TnI deficient conditions. To down-regulate TnI we used a validated RNAi (Casas-Tinto et al., 2016), genetically driven by the UAS/Gal4 binary system (Brand and Perrimon, 1993). We generated FLP/FRT clones using the heat shock inducible flipase (FLP) activity and *actin-Gal4* cassettes that allow identifying the TnI^RNAi^ expressing cells by GFP or RFP reporters (see **M&M**). In wing disc cells, Bazooka/Par-3 (Baz) is an apical polarity signal whereas Disc large (Dlg) is the baso-lateral counterpart (Kaplan et al., 2009; Roberts et al., 2012; Rolls et al., 2003). We monitored Baz by two different procedures, a GFP-tagged construct or an anti-Baz antibody. The data show that the depletion of *TnI* results in the delocalization of Baz as determined by either of the two visualizing procedures (**Fig. 2A,B**) (**Suppl. Fig. S1A, B**). Likewise, the basal polarity signal Dlg fails to accumulate at the basal domain in TnI depleted wing cells (**Fig. 2C,D**). This effect, however, is not bidirectional because depleting *Dlg* does not alter TnI localization (**Fig. 2E-G**). As a consequence of polarity loss after *TnI* depletion, cells undergo basal extrusion from the wing epithelium (see arrow in **Fig 2D**).

**Figure 2.**
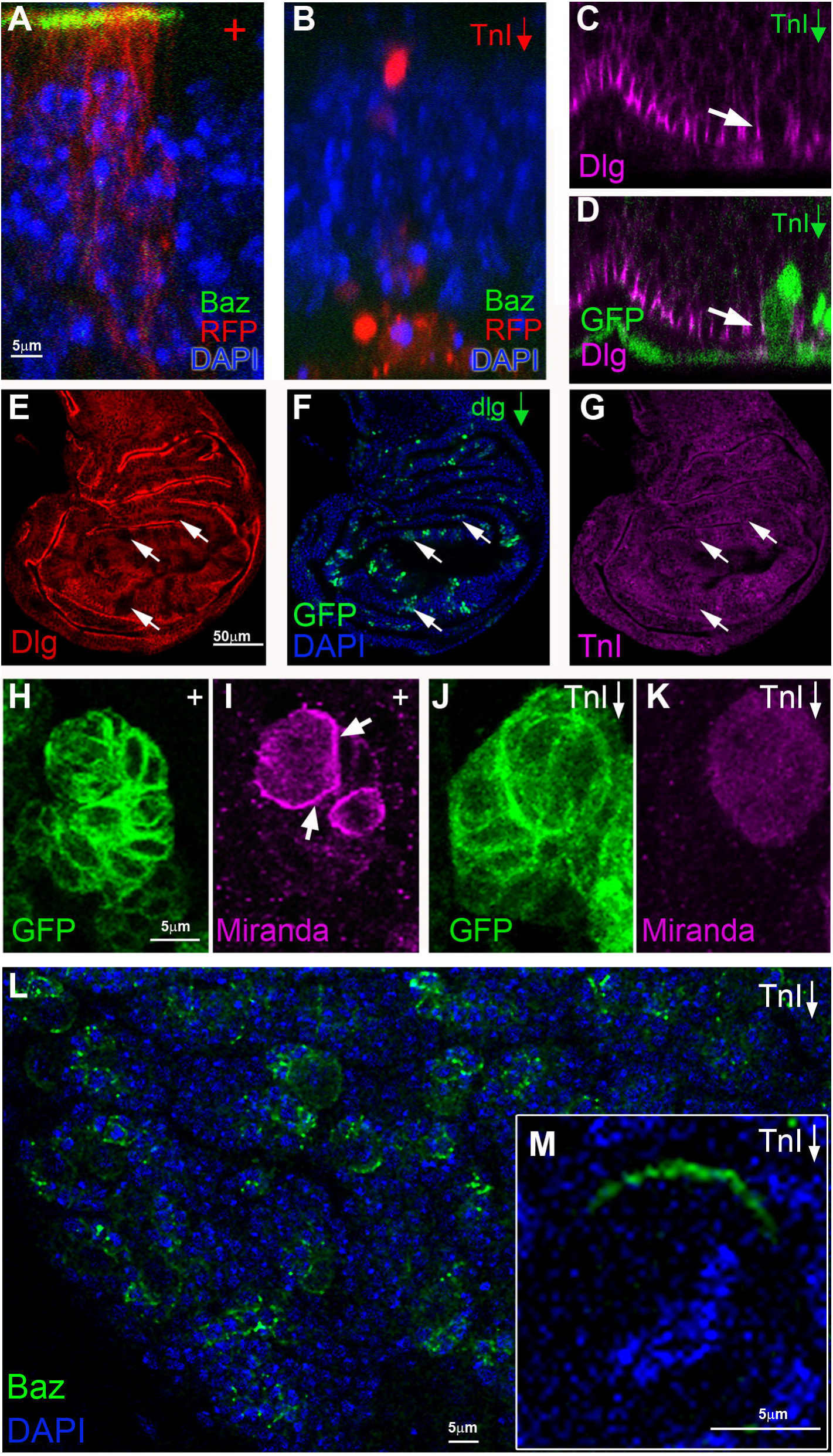
TnI interacts with polarity proteins. A-B) Z-axis view of a wing disc with an FRT/FLP clone (RFP) expressing Bazooka-GFP (**A**) or Bazooka-GFP plus *TnI*^*RNAi*^ (**B**). **C,D)** Z-axis view of a wing disc with TnI deficient cells (green) and immunostained for Dlg (magenta). Dlg mislocalizes in TnI deficient cells which are basally extruded from the epithelium (arrow). **E-G)** Wing disc immunostained for Dlg (red) and carrying GFP marked spots of *Dlg*^*RNAi*^ expressing cells (arrow heads) (green) show no effect on TnI expression (magenta). **H,I)** Larval neuroblasts (green) accumulate Miranda (magenta) in the basal pole (arrows in **I**) which is included in the derived ganglion mother cell. **J**,**K)** *TnI*^*RNAi*^ prevents polar accumulation of Miranda which remains homogenously distributed in the cytoplasm. **L,M)** *TnI*^*RNAi*^ does not interfere with polar accumulation of Baz in larval brain neuroblasts, magnification in **M**. Genotypes: *actin-FRT-Stop-FRT-Gal4/hs-FLP/UAS-RFP/UAS-TnI*^*RNAi*^*/UAS-Baz-GFP* (**A,B**); *actin-FRT-Stop-FRT-Gal4/hs-FLP/UAS-GFP/UAS-TnI*^*RNAi*^ (**C,D**) or *UAS-dlg*^*RNAi*^ (**E,F,G**); *worniu-Gal4/UAS-CD8-GFP/UAS-LacZ* (**H,I**); *worniu-Gal4/UAS-CD8-GFP/UAS-TnI*^*RNAi*^ (**J,K**) and *worniu-Gal4/UAS-TnI*^*RNAi*^*/UAS-Baz-GFP* **(L,M)**.

Miranda (Mir) is another basal polarity signal, albeit specific for neuroblasts. We addressed its localization in larval type II neuroblasts (*worniu-Gal4*) (**Fig. 2H,I**). Under *TnI* depletion, Mir remains homogeneously distributed in the cytoplasm (**Fig. 2J,K**). Thus, neuroblasts also require TnI to localize another basal polarity signal, Miranda in this case. Noticeably, the non-polar localization of Dlg in neuroblasts is not affected by *TnI* down-regulation, suggesting that the TnI-Dlg interaction is cell type specific (**Suppl. Fig. S1E, F**). Actually, Dlg is known to contribute to basal proteins targeting in neuroblasts through the interaction with lethal giant larvae (lgl) (Peng et al., 2000).

The contribution of TnI to localize AB polar factors, however, does not seem to be a general requirement for all polarity proteins. Scribbled (Scrib) expression and its basolateral localization in epithelial and salivary gland cells, or the apical accumulation of Baz in neuroblasts are not affected by *TnI* depletion (**Fig. 2L,M**) (**Suppl. Fig. S2A-F)**. These results reveal a degree of specificity in the mechanisms to localize polarity proteins that was unsuspected `hitherto`.

### TnI binds apico-basal polarity proteins

Co-immunoprecipitation assays (Co-IP) determined that TnI physically interacts with Baz as well as with Dlg. The interaction was validated in both directions; either pulling down from Dlg or from TnI (**Fig. 3A**). Since Baz and Dlg show opposite polar localizations, we determined if they form either a single or two independent complexes with TnI. To that end, we used triple Co-IP and antibody staining with anti-Baz, anti Dlg and anti-TnI. While Baz precipitated with TnI, no signal for Dlg was detected (**Fig. 3**). This result suggests that TnI forms independent complexes to position dorsal Baz and ventral Dlg. In addition to Baz, the apical complex includes atypical PKC (aPKC) (Harris and Peifer, 2005). We reproduced the Baz-aPKC binding in Co-IP assays (**Fig. 3B**). Furthermore, pulling-down from TnI also Co-IP aPKC and Baz (**Fig. 3B**). Since TnI is an actin-binding protein, these Co-IP data between TnI and selected AB polarity proteins indicate that their localization is mediated through actin filaments.

**Figure 3.**
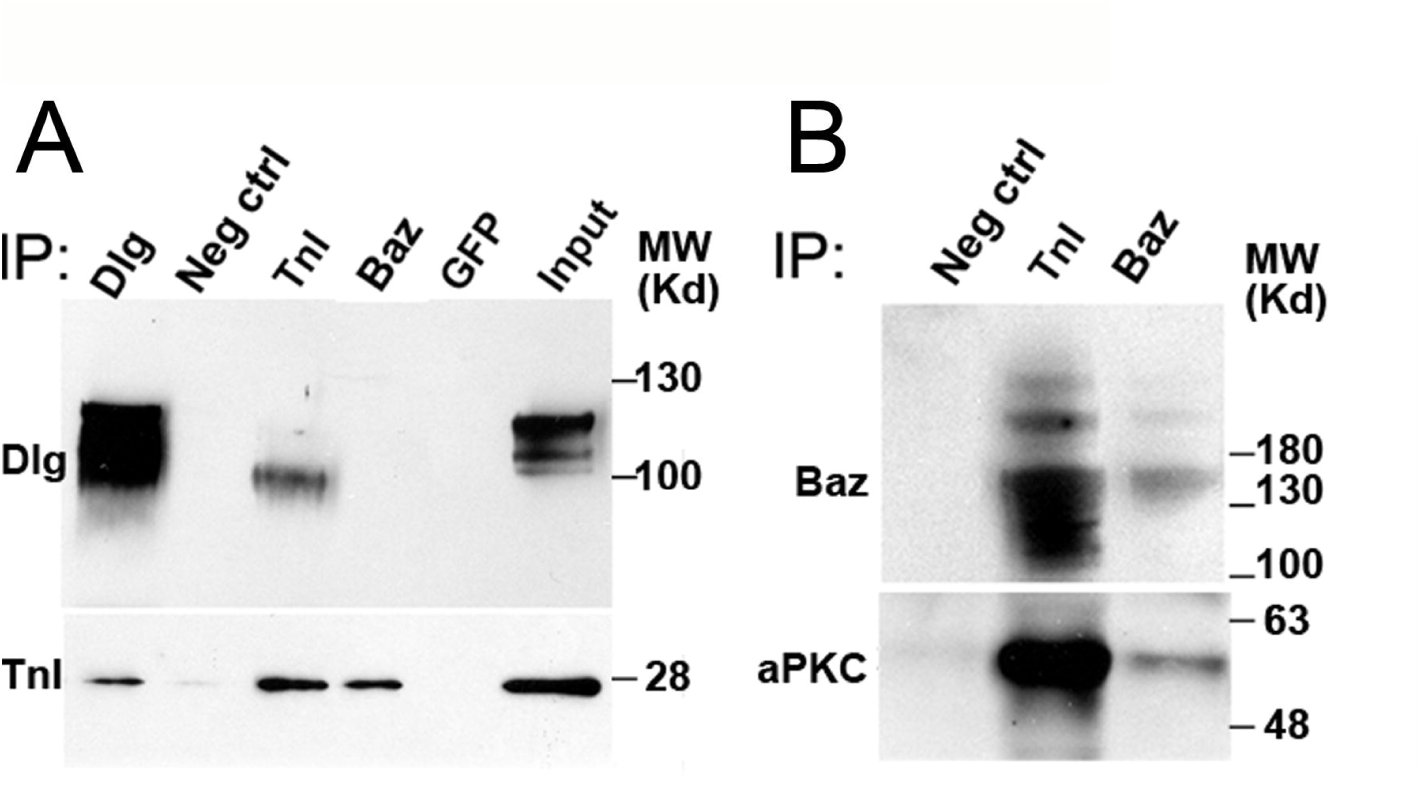
TnI co-immunoprecipitates with apico-basal polarity factors. A) Co-IP assays using antibodies against: Dlg, TnI, Baz or GFP (unrelated antibody used as negative control) and blotting against Dlg or TnI. Pulling-down Dlg brings TnI and *vice versa*, while pulling-down Baz brings TnI but not Dlg. **B)** Co-IP assays show that TnI brings down Baz and aPKC. Neg ctrl samples are beads with anti TnI without protein extract.

Apico-basal polarity complexes are linked to *adherens junctions* providing stability to the epithelium, among other functions (reviewed in (Tepass, 2012)). We monitored the integrity of these junctions, and the cytoskeleton in general, in TnI deficient wing cells using β-Catenin/Arm, E-Cadherin and γ-Tubulin as reporters, respectively. In these experiments, we generated large mosaics by driving the corresponding constructs to the whole posterior wing compartment (*en-Gal4*). The data show the expected reduction of reporter signals when TnI is down-regulated (**Fig. 4**). The weakening of these structures is expected to result in some form of cell unfitness leading to the observed extrusion from the epithelium by the neighboring cells (Baffet et al., 2012; Marinari et al., 2012).

**Figure 4.**
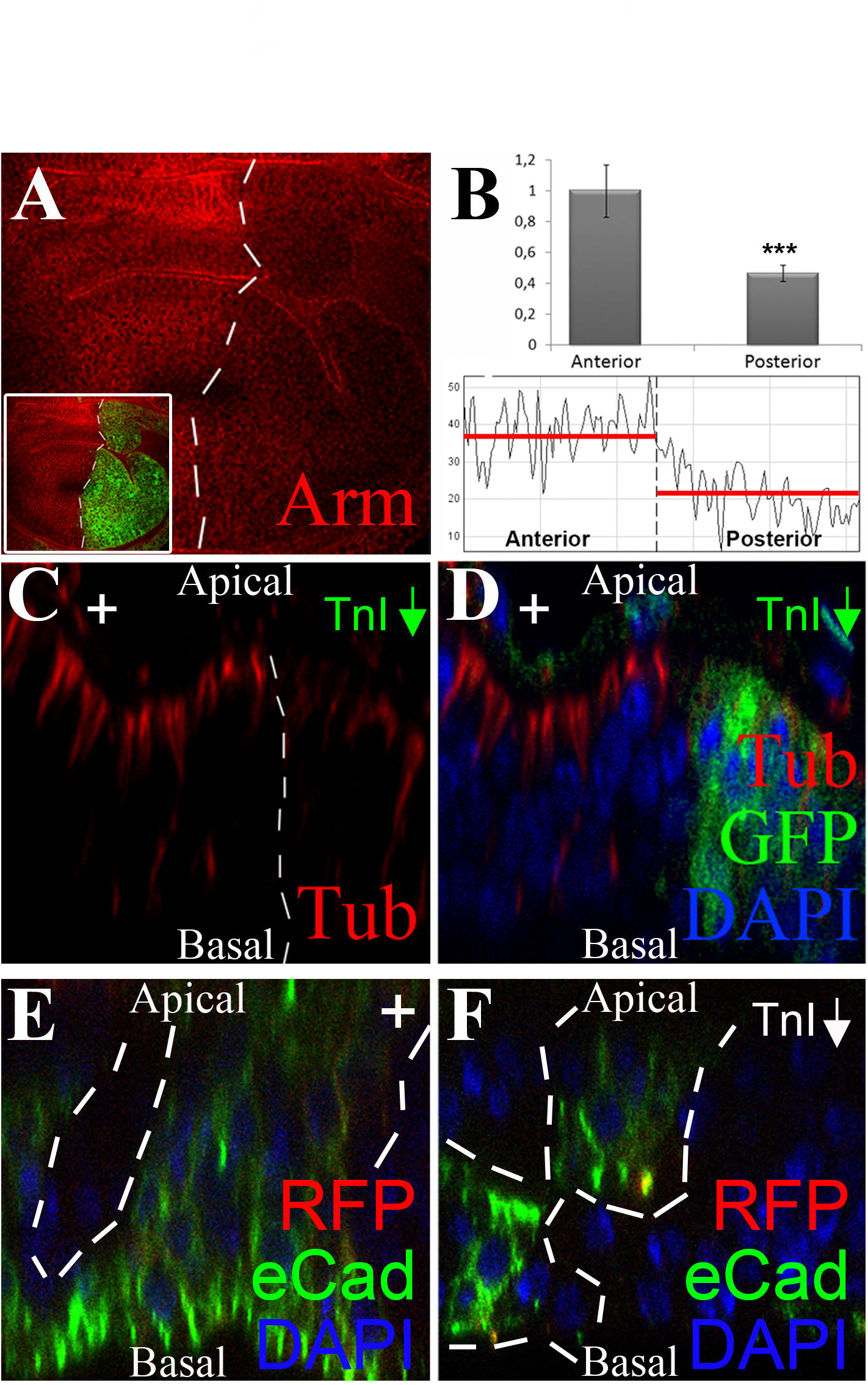
TnI depletion alters cytoskeletal structures. A) The down regulation of TnI in the posterior wing compartment (see inset) attenuates the immunosignal for β-Catenin/Armadillo (Arm). **B)** Quantification of the Arm immunosignal. Red dotted line indicates the average value in each wing compartment. **C,D)** Z-axis view of a wing disc immunostained against gamma-Tubulin (red) in which TnI has been depleted in the posterior compartment (green). Note the apical localization of gamma-Tubulin in the anterior compartment cells and its elimination in cells of the posterior compartment. **E)** Z-axis view of a wing disc expressing UAS-DE-Cadherin-GFP (green), to show its basal accumulation. **F)** Equivalent view of a disc with a clone of TnI depleted cells (dotted line). Note the mislocalization of DE-Cadherin. Genotypes: *en-Gal4/UAS-GFP/UAS-TnI*^*RNAi*^ (**A-D**); *hs-FLP/actin-FRT-Stop-FRT-Gal4/UAS-DE-CadGFP/UAS-RFP/UAS-TnI*^*RNAi*^ (**E,F**).

Searching for additional mechanisms involved in the localization of polarity factors, we noticed that mammalian PI3K mediates AB polarity in epithelial cells and contributes to basal membrane formation (Peng et al., 2015). Thus, we analyzed the potential effects of PI3K depletion in wing disc cells and found no alterations in the apical localization of TnI, Par3, or in the basal localization of Dlg (**Suppl. Fig. S2**). Nevertheless, like in mammals, fly PI3K must play a role in AB polarity because its upregulation rescues TnI deficient cells from apoptosis and extrusion due to polarity disruption (**Suppl. Fig. S3**). In addition, fly PI3K rescues TnI depleted cells from the effects on β-Catenin/Arm, γ-Tubulin and Dlg described above (**Suppl. Fig. S4**).

### TnI depletion is deleterious for the cell

Correct apico-basal polarity is required for normal cell proliferation in the wing disc. As we have shown previously, *TnI* depleted cells undergo cell competition and apoptosis after Caspase 3 activation, and FLP-out clones are eliminated from wing discs by 48 h AHS (Casas-Tinto et al., 2016).

Here, we aimed to analyze cell structural features caused by TnI depletion in the absence of cell death and elimination. To obtain that scenario, we monitored E-Cadherin or γ-Tubulin in *TnI* deficient cells under conditions in which cell competition is restrained by the up-regulation of survival signals or by the downregulation of cell death signals. The *death-regulator Need2-like caspase* (*Dronc*) encodes an endopeptidase involved in apoptosis (Yang et al., 2010). Its joint downregulation with TnI rescue cells from apoptosis but the levels of Armadillo are still reduced (**Fig. 5A-C**), as the TnI depletion alone does already (**Supp. Fig. S4**). The same effect was observed by upregulating the cell survival factor *secreted protein acidic rich cystein* (*Sparc*) (Portela et al., 2010) (**Fig. 5D-F**) or with the down regulation of the cell death signal *flower* (*fwe*) (Rhiner et al., 2010) (**Fig. 5G-I**). Concerning Tubulin, cells rescued from apoptosis by the overexpression of p35 (Hay et al., 1994) still show the γ-Tubulin loss from the apical end that TnI depletion causes by itself (**Fig. 5J-L**). In summary, TnI loss causes defects in epithelial cell adhesion and tubulin cytoskeleton that are maintained even if cells are rescued from apoptosis and elimination. This suboptimal status of imaginal wing cells is likely to cause the morphological defects observed in the resulting adult wings (**Fig. 5M-P**).

**Figure 5.**
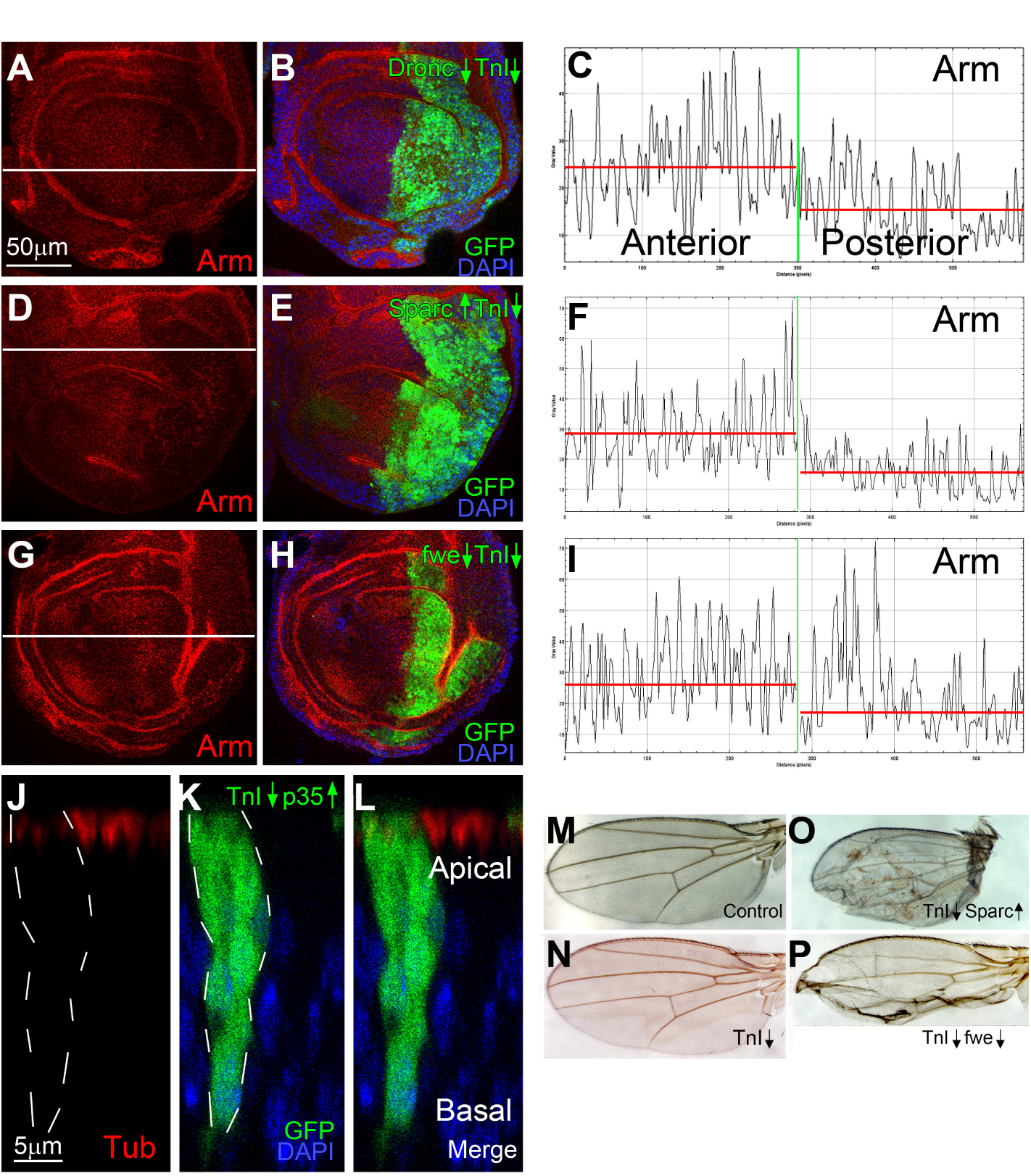
Cytoskeletal defects due to TnI depletion can be separated from apoptosis. A-I) The down regulation of TnI in the posterior wing compartment attenuates the immunosignal for Armadillo even if cell death is prevented by down-regulating *Dronc* (**A,B**) or up-regulating *Sparc* (**D-F**) or *fwe* (**G-I**). Digital quantification of Arm signal (**C,I,F**) corresponds to the white line in **A,D** and **G**. The red line indicates the average value in each wing compartment. **J-L**) Z-axis view of a wing disc clone of TnI depleted cells rescued from apoptosis by up-regulating p35 (green) showing the attenuation of γ-tubulin immunosignal (red). **M-P)** Adult wings from larvae expressing multiple clones of TnI depleted cells rescued from apoptosis by *Sparc* or *fwe*. Genotypes: *en-Gal4/UAS-TnI*^*RNAi*^ plus *UAS-Dronc*^*RNAi*^ or *UAS-Sparc* or *UAS-fweLoseA/B*^*RNAi*^ (**A-I**); *hs-FLP/actin-FRT-Stop-FRT-Gal4/UAS-GFP/UAS-TnI*^*RNAi*^*/UAS-p35* (**J-L**); *hs-FLP/actin-FRT-Stop-FRT-Gal4/UAS-GFP/UAS-TnI*^*RNAi*^*/ UAS-Sparc* or *UAS-fweLoseA/B*^*RNAi*^ (**M-P**).

E-Cadherin and γ-tubulin are also relevant for mitotic spindle orientation and chromosome segregation (Baffet et al., 2012; Wang et al., 2004). In that context, we monitored the cell centrosome using a GFP-tagged form of *asterless* (*pEYFP.asl*^*FL*^) that binds γ-Tubulin in the pericentriolar material of the centrosome (Gopalakrishnan et al., 2011). *TnI* mutants show aberrant number of centrosomes around single nuclei in the syncytial embryo stage (**Fig. 6A,B**). The feature is also reproduced in wing disc cells expressing a *TnI*^*RNAi*^ at 24-36 h post clone induction, a time prior to cell extrusion from the epithelium (**Fig. 6C-G**) confirming that both tools to deplete TnI, RNAi and mutants, yield the same phenotype and are similarly effective.

**Figure 6.**
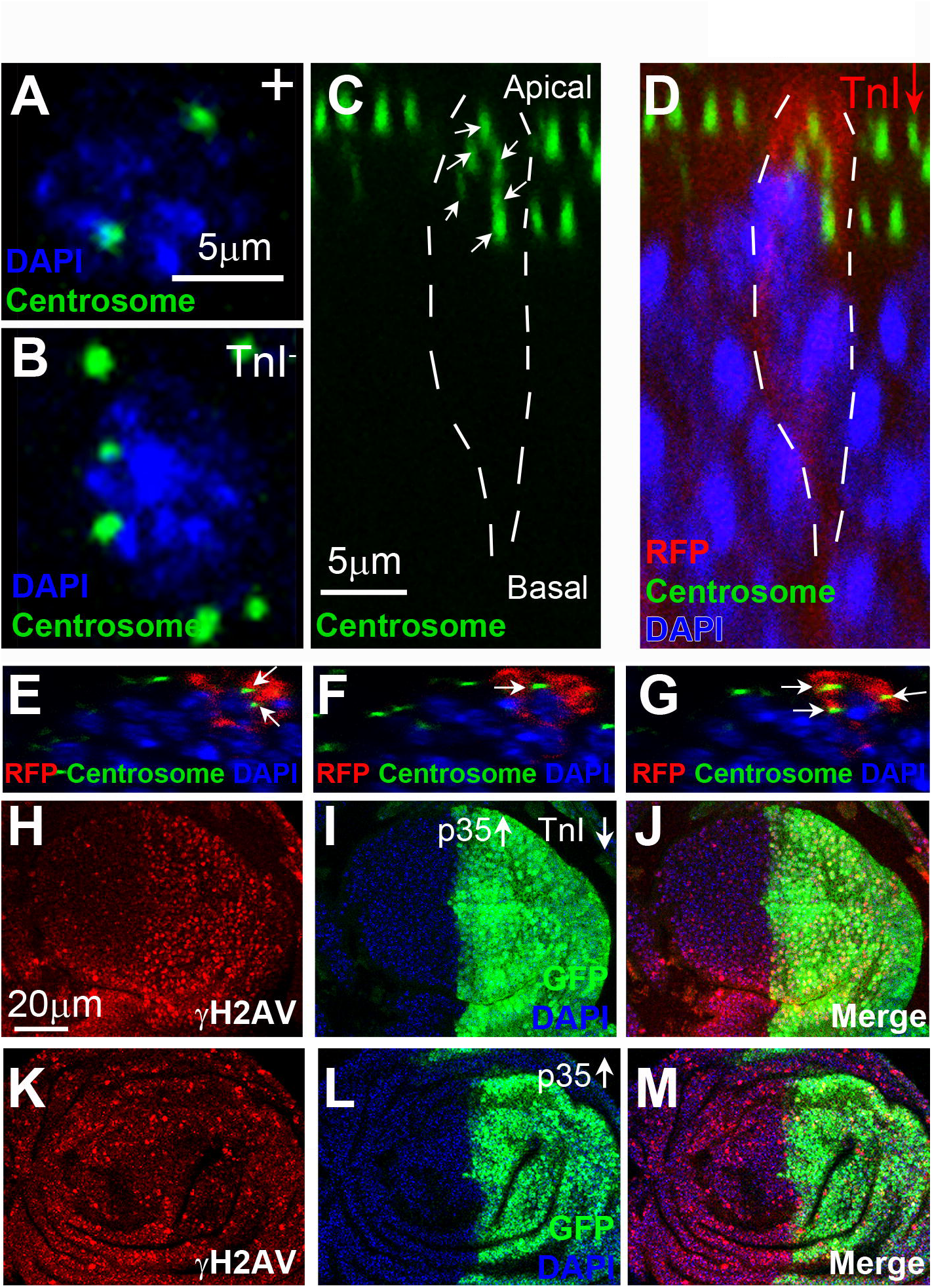
TnI is required for normal mitosis and chromosome integrity. A,B) Normal syncytial embryo nuclei exhibit two asters (**A**) but null TnI embryos often show aberrant number of asters (**B**). DAPI (blue) marks DNA and *pEYFP.asl*^*FL*^ (green) marks asters. **C-G)** Z-axis view of a wing disc clone of TnI depleted cells. Note the additional asters (arrows). **H-J)** TnI depletion in the posterior wing compartment (green) rescued from apoptosis by p35 increases genome instability as revealed by anti-γH2AV antibody (red). **K-M)** The expression of p35 alone, however, does not increase the anti-γH2AV signal. Genotypes: *pEYFP.asl*^*FL*^ (**A**) and *pEYFP.asl*^*FL*^*TnI*^*23437*^ (**B**) embryos; *hsFLP/pEYFP.asl*^*FL*^, *actin-FRT-Stop-FRT-Gal4 UAS-RFP/UAS-TnI*^*RNAi*^ (**C-G**); *en-Gal4 UAS-GFP/ UAS-TnI*^*RNAi*^*/UAS-p35* **(H-J)**; *en-Gal4 UAS-GFP/UAS-p35* **(K-M)**.

### TnI depleted cells rescued from apoptosis still show genome instability

Chromosome mechanics is likely affected if centrosomes are abnormal. Previous reports indicate that microtubules and conserved PAR proteins are essential mediators of cell polarity, and mitotic spindle positioning depends on heterotrimeric G protein signaling and the microtubule motor protein dynein (Ahringer, 2003). In particular, a Baz-centrosome positive feedback loop contributes to the maintenance of the *adherens junctions* and cell integrity (Jiang et al., 2015). To monitor the genomic stability, we relied on the γH2AV antibody. This histone H2 reporter is a standard indicator of genome integrity (Madigan et al., 2002). As in previous experiments, apoptosis was prevented by up-regulating p35. The data show a significant increase of the γH2AV signal in the TnI deficient cells (**Fig. 6H-J**). As control, cells with up-regulated p35 alone did not show changes in γH2AV signal (**Fig. 6K-M**). Two additional methods to rescue TnI depleted cells from apoptosis, downregulation of *fwe* or upregulation of *Sparc*, also maintained the genome instability (**Suppl. Fig. S5**). It is worth noting that cells exhibiting genome instability and rescued from apoptosis are still able to proliferate. This is consistent with equivalent experiments in which genome instability had been elicited by alternative methods such as mutations in genes involved in the spindle assembly checkpoint (*bub3* and *rod*), spindle assembly (*asp*), chromatin condensation (*orc2*), or cytokinesis (*dia*) (Dekanty et al., 2012).

In summary, the events triggered by TnI depletion follow the order: apico-basal cell polarity loss, structural abnormalities in *adherens junctions* and cytoskeleton, genome instability and final apoptosis. The last step can be prevented allowing proliferation to some extent, but cells acquire a suboptimal state which is prone to yield morphological abnormalities after differentiation. In addition, genome instability seems mechanistically different from the rest of AB polarity related events.

## Discussion

This study shows that TnI contributes to the AB cell polarity and binds directly certain proteins which can accumulate in the apical or in the basolateral region of the cell. The binding partners for TnI, however, are selective according to the cell type. Thus, the basal Dlg of epithelial cells requires TnI for its preferential localization, but it is not perturbed if TnI is depleted in neuroblasts where Dlg is not polar. These features unveil a diversity of mechanisms to allocate specific proteins to polar domains that was unsuspected hitherto. In addition to the previously known AB polarity factors, TnI is revealed as an apical factor at least in epithelial cells. Preliminary observations in salivary gland cells seem to indicate a still unexplored apical localization within the cell nucleus.

The mechanism by which TnI allocate apical (Baz/Par3) and basolateral (Dlg) factors in epithelial cells includes their direct binding, albeit in different complexes. That is, TnI would associate with different partners according to the cargo destination and cell type. In essence, TnI is an actin-binding protein and this feature is likely to sustain the force needed to move polarity factors within a cell. Also, the contribution of TnI helps to explain the regulation of force from the polarity complexes with the neighboring cells, through the adherence junctions that these complexes help to establish (Wang et al., 2012). Similar to its role during muscle sarcomere contraction, TnI would regulate actin filaments in their sliding along, most likely, a membrane-anchored isoform of myosin. Thus, actin-based movements in a cell seem to use the same basic mechanism as in muscles. The role of PI3K in the TnI mediated mechanisms for AB polarity seems to be different between vertebrates and *Drosophila*, although this kinase clearly participates in the fly’s AB cell polarity mediated by TnI. The phosphorylated substrate of PI3K in this context remains to be identified. However, a particular form of myosin, myosin-1s, has been reported to target membrane phosphoinositides in addition to F-actin (Rajendraprasad et al., 2018). This scenario would be suitable to integrate PI3K, TnI, F-actin and myosin into a force generating complex to move AB polarity factors.

TnI depleted cells lose AB polarity and reduce proliferation. Thus, TnI represents a link between these two cellular properties. TnI would regulate actin fibers that control cell shape. Loss of polarity and reduced proliferation cause some form of unfitness with deleterious effects. In wing discs, cell unfitness results in apoptosis and extrusion. One of the proposed mechanisms for wing cells extrusion is dependent on actin/myosin (Rosenblatt et al., 2001; Slattum et al., 2009). In RasV12 transformed epithelial cells, extrusion is mediated by a mechanism that depends on EPLIN/LIMA1 and that activates myosin II and PKA (Ohoka et al., 2015). Cell competition as defined in *Drosophila* is considered to maintain organ shape by extrusion of the so called loser cells. The probability of loser cell extrusion correlates with the extent of surface interaction with the surrounding winner cells (Levayer et al., 2015). The process, however, is likely to be more heterogeneous than initially thought, since cell elimination can also be triggered away from the loser/winner interface, by means of cell crowding, a phenomenon named “super-competition” (Levayer et al., 2016). Elimination of unfitted cells is also observed in vertebrates, particularly in epithelia, where it has been renamed as “Epithelial-defence-against-cancer, EDAC” (Kajita and Fujita, 2015).

The cell phenotypes reported here are compatible with a causal link between loss of AB polarity, cell competition, extrusion or apoptosis. An additional feature, genome instability, has been proposed to be part of this sequence of events; in particular to explain genetic heterogeneity in tumor metastases (reviewed in (McGranahan and Swanton, 2017). In this study on TnI, we have attempted to determine if genome instability is indeed a consequence of AB polarity loss, at least the polarity loss mediated by TnI. Experiments in which TnI depleted cells are prevented from extrusion or apoptosis by means of protecting them with p35, *Sparc* or *fwe*, clearly indicate that genome instability is a process mechanistically different from cell competition due to cell polarity loss, and it does not lead to cell death necessarily. Further, cells exhibiting genome instability are able to proliferate to some extent. Actually, this feature is in agreement with other experiments in which genome instability was triggered by other methods (Dekanty et al., 2012).

The fly’s apical complex is conserved in vertebrates (Harris and Peifer, 2005; Zhu et al., 2011) and cell polarity genes are also tumour suppressors (Hanada et al., 2000; Humbert et al., 2008; Iden et al., 2012; Zhu et al., 2014). As shown previously, TnI enhances the oncogenic properties of *lgl*, *N*, and *Ras* in *Drosophila* and it is at the origin of certain types of cancers in humans (Casas-Tinto et al., 2016). Thus, the functional link between AB polarity and proliferation through TnI is expected to be conserved as well. Given the selectivity of the TnI contribution to AB polarity and its requirement for cell proliferation, it may become a suitable target for cell type specific therapies.

## Materials and Methods

### Fly strains and genetic crosses

The fly stocks used are from the Bloomington Stock Center (Fly Base) except as indicated. As null TnI mutation we used a rearrangement located in the regulatory region of the *wupA* gene, *Df(1) TnI*^*23437*^ (Marin et al., 2004). To induce TnI loss-of-function cell mosaics we used the Bloomington line *P{TRiP.JF02172}attP2* (BL#31893) following a comparative study with other available RNAi lines(Casas-Tinto et al., 2016). Constructs *UAS-p35*, *UAS-fweLoseA/B*^*RNAi*^ and *UAS-Sparc* were obtained from E. Moreno (Portela et al., 2010; Rhiner et al., 2010), UAS-DE-Cadherin-GFP (Gorfinkiel and Arias, 2007) and *w; worniu-Gal4; Ase-Gal80* were from C. González. Line *pEYFP.asl*^*FL*^ marks the centriole specific Spindle abnormal assembly protein 6 and was obtained from C. González (Varmark et al., 2007). FLP/FRT mosaics were obtained by heat shock of larvae carrying the constructs *hs-FLP*; *actin-FRT-yellow-Stop-FRT-Gal4,UAS-GFP/+* combined with the desired *UAS-x* construct. The expression of the flipase after a heat shock (AHS) excised the FRT-flanked “*yellow*” cassette and generated Gal4-expressing clones. These clones overexpress the respective UAS-constructs driven by the ubiquitous *actin* promoter (*act > Gal4*). Two versions of this cassette were used, one including *UAS-GFP* and the other including *UAS-RFP*. We used a GFP-Scrib fusion protein to visualize Scrib together with a RFP-histone reporter to visualize nuclei: scribbled-GFP his1Av-mRFP.

### Mosaic generation and clone size measurement

Crosses were set with 10 females and 10 males per vial at 25°C and vials were changed every 72 hours to avoid overcrowding. Mitotic recombination clones were obtained by delivering a heat shock (1hour at 37°C) during 2^nd^ instar larvae. 48 or 72 hours after heat shock 3^rd^ instar wandering larvae were dissected. FLP-out clones were obtained by identical protocol but with different heat shock conditions (8 minutes at 37°C). Control cultures were run in parallel. Wandering LIII larvae were dissected for clone screening.

### Immunohistochemistry and Immunoprecipitation

Tissue samples were fixed in formaldehyde 4% 25 min and stained according to standard protocols. Antibodies: anti-Active caspase3 (1/100 Cell Signaling), anti-γH2AV (1/100 DSHB), anti-Lamin (DSHB 1/100), anti-Actin (DSHB), anti-Tub (1/100 SIGMA), anti-Dlg (1/20 DSHB), anti-Par3 (1/50 DSHB), anti-Crumbs (1/100 DSHB), anti-Scrib (1/20 Santa Cruz Biotechnology) anti-Arm (DSHB 1/100), anti-miranda (1/100) gift from Rita Sousa Nunes and anti-β-galactosidase (1/50 DSHB). Fluorescent secondary antibodies Alexa 488, 569 and 647 were used (1/200 Invitrogen). All images were obtained with a LEICA TCS-SP5 confocal microscope and processed with the software-assisted Bitplane’s Imaris for measurements. The SC1 mouse monoclonal Troponin-I antibody was raised against the peptide PDGDPSKFAS (Abmart Inc.) (Casas-Tinto et al., 2016). TnI-Dlg co-immunoprecipitation was detected in whole fly extracts using specific mouse SC1 anti-TnI (1/100), mouse anti-Dlg (1/10), mouse anti-Baz (Par3) (1/100) (DSHB (1/50), rabbit anti-aPKC (1/100) and mouse anti-GFP (1/100) (Roche) as a negative control, coupled to G-Sepharose beads (GE healthcare). Pull-down assays were done with Canton-S strain protein extracts. Antibody immobilization to Sepharose beads was performed for 8 h at 4°C and subsequent incubation with protein extracts was conducted in the presence of 0.25% BSA for 2 hours at RT. Immunoprecipitated proteins were subjected to Western blot developed with True Blot secondary antibodies (eBioscience). Negative controls (Neg Ctrl) are G-Sepharose beads without primary antibodies. Input is 1/10 of the original protein extract.

### Western blotting

Western blots were obtained by standard procedures (Invitrogen) and protein bands were quantified by densitometry using ImageJ software with the “GelAnalyzer” option (See further details in http://www.di.uq.edu.au/sparqimagejblots).

### Statistics

The area of GFP positive clones was quantified using Image-J A 1.44a and Imaris (Bitplane) software. Averages and standard deviations (SD) were calculated from the ratio GFP/DAPI area. Statistical significance was calculated with the two-tailed Student’s *t*-test. Significance levels are indicated as * p<0,05**p<0,005***p<0,001. Number of samples N>8 animals in all cases.

## Acknowledgements

We appreciate fly strains from Bloomington Stock Center. Research was funded by grants BFU2012-38191 and BFU2015-65685-P from the Spanish Ministry of Economy.

## Supplementary Figure Legends

**Figure S1.-TnI is required for the polar localization of certain polarity factors. A-B)** Z-section of wing disc with an RFP marked clone expressing TnI^RNAi^ and stained with an anti-Baz antibody. Note the cell autonomous removal of Baz (green) in the TnI down-regulated cell (red). **C,D)** Z-section of wing discs with clones (GFP) of cells expressing RNAi for *aPKC* and immunostained against TnI (red). The TnI signal is not affected. **E,F)** The non-polar localization of Dlg (red) is not affected by the down-regulation of TnI in type II neuroblasts and their descendants. Genotype: *worniu-Gal4; UAS-TnI*^*RNAi*^; *UAS-GFP*. **G,H)** Salivary gland cells with TnI depleted clones (red), induced in a background that constitutively expresses GFP-tagged Scrib. Cell nuclei also express His-1Av-mRFP (genotype: *hsFLP/+; actin-FRT-Stop-FRT-Gal4UAS-RFP/Scrib-GFP, his1Av-mRFP; UAS-TnI*^*RNAi*^*/+*). The basolateral polarity factor Scrib is not altered by the loss of TnI. **I,J)** Wing disc clones generated by mitotic recombination (genotype: *Df(1) TnI*^*23437*^ *FRT18A/y w ubi-GFP FRT18A ; hsFLP38/+*) stained with anti-Scrib (red). Larvae were subjected to a 45-min heat shock at 37°C and harvested 48h later. The TnI null clone (arrow) is surrounded by the twin (dotted line) (intense green) clone. The TnI clone is very small due to cell competition and extrusion. The localization of Scrib, revealed by a mouse-antibody, remains normal in all wing cells.

**Figure S2.-PI3K does not alter AB polarity factors in wing epithelial cells. A-C)** Z-axis view of a wing disc showing a clone of PI3K depleted cells (red) and immunostained against TnI. **D-F)** Equivalent case for a disc stained with anti-Par3. **G-I)** Equivalent case for a disc stained with anti-Dlg. None of these apical (TnI, Par3) or basal (Dlg) factors are affected by the depletion of PI3K.

**Figure S3.-PI3K does rescue TnI depleted cells from apoptosis and extrusion.** Wing disc with a large clone of TnI depleted cells that co-express PI3K resulting in their rescue from apoptosis and extrusion. Z-axis views shown in inlets correspond to the dotted white lines. Genotype: *actin-FRT-Stop-FRT-Gal4/hs-FLP/UAS-GFP/UAS-TnI*^*RNAi*^*/UAS-PI3K.*

**Figure S4.-PI3K does rescue TnI depleted cells from cytoskeletal defects. A-C)** Z-axis view of a wing disc with TnI depleted/PI3K upregulated clones (green) immunostained against Arm (red). Note that the apical localization of Arm remains as normal. **D-F)** Equivalent case stained with anti-γ-Tubulin (red). Note that the apical accumulation of γ-Tubulin is not altered. **G-I)** Equivalent case stained with anti-Dlg (red). Note that the basal localization of Dlg remains unaffected in spite of the TnI depletion. Genotype: *actin-FRT-Stop-FRT-Gal4/hs-FLP/UAS-GFP/UAS-TnI*^*RNAi*^*/UAS-PI3K.*

**Figure S5.-The rescue of TnI depleted cells from apoptosis does not prevent genome instability. A,B)** Rescue of TnI depleted cells by *flower (fwe)* knockdown in the posterior compartment. **C,D)** Equivalent rescue by *Sparc* overexpression. **E)** Quantification of H2AV immune positive spots in the previous genotypes. Note that genome instability, as monitored by anti-γH2AV, persists.

